# Human mesenchymal stem cells and derived extracellular vesicles reduce sensory neuron hyperexcitability and pain-related behaviors in a mouse model of osteoarthritis

**DOI:** 10.1101/2022.02.25.478196

**Authors:** Minji Ai, William E. Hotham, Luke A. Pattison, Qingxi Ma, Frances M.D. Henson, Ewan St. John. Smith

## Abstract

Osteoarthritis (OA) is a common degenerative joint disease characterized by joint pain and stiffness. In humans, mesenchymal stem cells (MSCs) and derived extracellular vesicles (MSC-EVs) have been reported to alleviate pain in knee OA. Here, we used the destabilization of the medial meniscus (DMM) mouse model of OA to investigate mechanisms by which MSCs and MSC-EVs influence pain-related behavior. We found that MSC and MSC-EV treated DMM mice displayed improved OA pain-related behavior (i.e. locomotion, digging and sleep) compared to untreated DMM mice. Improved behavior was not the result of reduced joint damage, but rather knee-innervating sensory neurons from MSC and MSC-EV treated mice did not display the hyperexcitability observed in untreated DMM mice. Furthermore, we found that MSC-EVs normalize sensory neuron hyperexcitability induced by nerve growth factor *in vitro*. Our study suggests that MSCs and MSC-EVs may reduce pain in OA by direct action on peripheral sensory neurons.

**Teaser:** Mesenchymal stem cells and secreted extracellular vesicles normalize sensory neuron excitability to reduce pain.

## Introduction

Osteoarthritis (OA) is a debilitating musculoskeletal disease affecting over 250 million people worldwide (*1*). Chronic pain is the primary OA symptom and the major driver for both seeking medical attention and clinical decision making (*2*, *3*). Poorly managed OA pain can lead to limited joint function (*4*), reduced quality of life (e.g. compromised sleep quality, anxiety, and depression) (*5*, *6*), and disability in patients (*7*). Unfortunately, currently used pharmacological treatments for OA pain (e.g. non-steroidal anti-inflammatory drugs and opioids) fail to provide sufficient pain relief and are often associated with unwanted side effects following long-term use (*8*). Thus, managing OA pain remains challenging and requires disease specific analgesics to address this unmet clinical need.

Peripheral input is a major contributor to OA pain as demonstrated by reduced pain in OA patients following: i) intra-articular injections of the local anesthetic lidocaine (*9*), ii) a peripherally restricted anti-nerve growth factor (NGF) antibody (*10*) and iii) total knee replacement (although pain persists in some patients) (*11*). Moreover, in rodents, inhibition of nociceptor activity with the quaternary anesthetic QX-314 ameliorated early OA pain (*12*), and we have previously shown that pain behaviors following joint injury can be reversed through chemogenetic inhibition of knee-innervating sensory neurons (*13*). Furthermore, in the monoiodoacetate model of OA in rats, it has been shown that knee-innervating extracellular electrophysiological recordings become sensitized early after disease onset (from day 3) and that this is maintained, whereas bone-innervating afferents only become sensitized late in disease (day 28) (*14*). The OA joint contains multiple cell types and mediators that have been identified as drivers of OA pain. Studies have identified several key molecules that are thought to drive OA pain and thus have been developed as disease specific pain target. For example, NGF was first identified as a pain target for OA as its expression was elevated in a murine OA model (*15*) and treatment with soluble NGF receptor tropomyosin receptor kinase A (TrkA) (*15*), anti-NGF antibody (*16*), and inhibition of the TrkA receptor (*17*) can all effectively suppress pain like behavior in rodent OA models. Moreover, a number of anti-NGF antibodies have demonstrated clinical efficacy in managing OA pain in patients, but the risk of causing rapidly progressive OA (perhaps in part by removing the protective effect of reduced weigh bearing on the diseased joint) has thus far prevented their clinical application (*18*). In addition, there has also been significant interest in the chemokine CCL2: animal studies revealed that blockade of the CCL2 receptor CCR2 improves pain symptoms in murine OA (*19*), and absence of both CCL2 and CCR2 delay OA pain development (*20*). Similarly, there is gathering evidence for a role of the aggrecan 32-mer fragment activating Toll-like receptor 2 to drive OA joint pain (*21*).

In search of a mechanism based therapeutic for OA, mesenchymal stem/stromal cell (MSC) therapy has emerged as a promising treatment, with clinical trials demonstrating pain relief and improved joint function in OA patients (*22*). The typical OA joint is characterized by cartilage loss and synovitis, which can be improved by MSCs primarily through immunomodulation. MSCs exert a strong immunomodulatory effect through the secretion of soluble factors such as anti-inflammatory proteins (e.g. Tumour necrosis factor (TNF)-α–stimulated gene 6 protein (TSG-6) (*23*)) and growth factors (e.g. transforming growth factor beta (TGF-β) (*24*)), which lead to analgesic and anti-catabolic effects in OA joints (*25*). The effects of MSCs, then, are to improve the joint microenvironment. However, a further possibility exists that they may directly alter the nociceptive input, which would contribute to the pain relief experienced by those with OA. However, a direct link between MSCs and nociception in OA remains unexplored, i.e., do MSCs affect neuronal excitability?

Despite promising outcomes, the clinical use of MSCs faces a number of safety concerns such as potential tumorigenicity (*26*). Therefore, extracellular vesicles (EVs) secreted by MSCs, have been proposed as an alternative to MSCs for treating OA, indeed, increasing evidence has attributed the therapeutic effects of MSCs to their paracrine secretion, especially of EVs (*27*–*29*). EVs are small sized, membrane bound vesicles (30–200 nm) that are secreted into the extracellular space by cells, including MSCs (*30*). Within EVs, there is a rich profile of biomolecules, including proteins, lipids, and nucleic acids, which have strong immunomodulating and chondroprotective properties (*31*). Although MSC derived EVs (MSC-EVs) are a highly heterogenous population, they can be broadly distinguished into three types based on their biological origins: exosomes, microvesicles and apoptotic bodies (*32*). Exosomes are small vesicles are secreted through a fusion of endosomal multi-vesicular bodies (MVBs) with the plasma membrane (exosomes, 30–120 nm) (*33*), while microvesicles are formed through the direct outward budding of cell membrane (microvesicles, 100–1000 nm) (*34*). Preclinical studies show that MSC-EVs derived from various sources (e.g. adipose, bone marrow, and umbilical cord MSCs) exert a similar therapeutic effect to their source cells in different OA models, such as inhibiting joint inflammation and promoting cartilage repair (*35*). However, the analgesic effects of MSC-EVs in OA remains unknown. In the present study, we aimed to determine to what extent either MSCs or MSC-EVs provide analgesia through studying their impact on nociception in the OA joint. We hypothesized that MSCs and MSC-EVs would improve OA pain via direct modulation of sensory neurons innervating the joint.

## Results

To test the hypothesis that MSCs and MSC-EVs directly modulate joint-innervating neurons to produce pain relief, we surgically induced knee OA in 10-week-old, male C57Bl6/J mice by conducting destabilization of the medial meniscus (DMM) surgery and randomly assigned mice into 4 experimental groups: sham, DMM, DMM+MSCs, DMM+MSC-EVs (Fig. 1A). Human MSCs were purchased commercially (Lonza, UK) and derived MSC-EVs were harvested and characterized as previously described (Fig.S1) (*36*). To exclude the regenerative effects of MSCs and MSC-EVs in OA that can be observed when administered at week 4 post DMM surgery (*37*), we started MSC/MSC-EV treatment from 12 weeks post-DMM surgery at which point OA is well established (Fig. 1A).

**Fig. 1.**
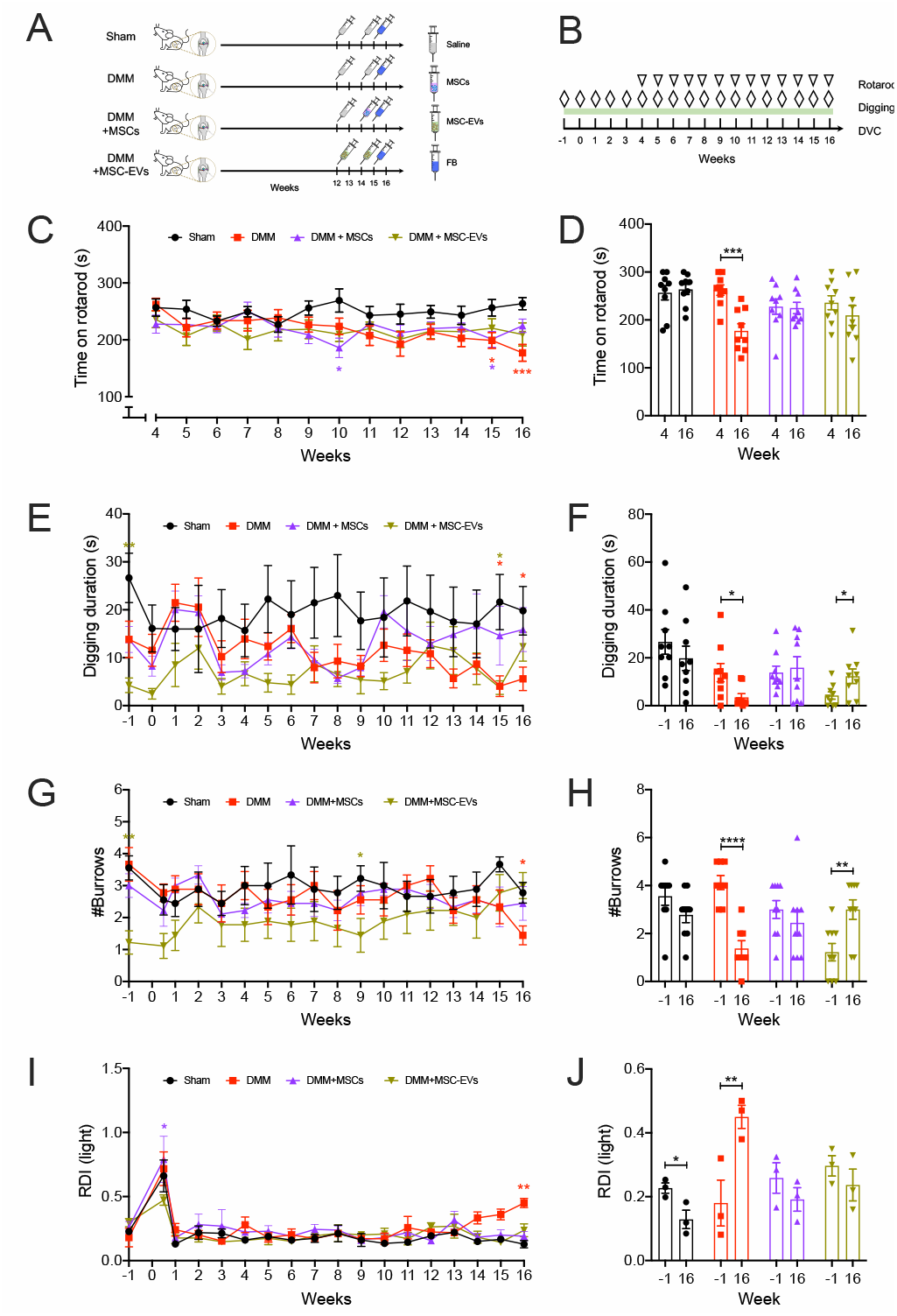
MSCs and MSC-EVs improves knee joint pain related behavior change in DMM mice. (A) Schematic experimental design of in vivo study (n=9/group). (B) Timeline of conducted behavior tests. Total time mice spend on the rod (C) at each week, and comparison of time on rod within each mouse group at week 4 and week 16 post-surgery. The total time mice spend digging during the testing period (E) at different weeks, and the comparison of digging duration at pre-surgery and at week 16 post-surgery within each mouse group. The number of burrows mouse dug by mice at the end of each test (F) at each week, and comparison of burrows dug at pre-surgery and at week 16 post-surgery with each mouse group (H). Light period RDI value for mice during experimental period (I) and comparison of light period RDI at pre-surgery and at 16-week post-surgery with each mouse group (J). *p<0.05, **p<0.01, ***p<0.001, **** p<0.0001. ns, no significant difference. Two-way ANOVA with Dunnett’s multiple comparisons test was used for behavior changes among four experimental groups across time series (C, E, J, I). Unpaired t test was used to compare behavior values at two different time points within each mouse group (D, F, H, J).

### MSCs and MSC-EVs improve pain-related behavior changes in DMM mice

To examine if MSC or MSC-EV treatment improves pain-related behavior in DMM mice, we used three different methods to monitor mouse behavior: rotarod test, digging test, and Digital Ventilated Cage^®^(DVC) system. All these measurements examine how DMM-induced pain affects normal mouse behavior, rather than evoked pain, to better align with how on-going pain affects the behavior of those individuals living with OA pain. Because the rotarod forces an animal to behave in a certain way and ability to perform is likely to be impacted by surgery, it was only conducted weekly from week 4 post-surgery, whereas the digging test was carried out weekly from one week pre-surgery and DVC measurements were made for the duration of the study, also from one week pre-surgery (Fig. 1B).

The daily use of a painful joint lead to behavioral adaptation affecting gait resulting in a locomotion deficit (*38*). Previous studies reported reduced locomotion in DMM mice after 16 weeks using rotarod tests (*39*, *40*). We observed that untreated DMM operated mice started to spend significantly less time on the rotarod than sham mice at week 15 (week 15: Sham: 256.4 ± 14.6 sec vs. DMM: 199 ± 13.88 sec; p = 0.03) and at week 16 (week 16: Sham: 256.4 ± 14.6 sec vs. DMM: 177.1 ± 14.77 sec; p = 0.0008, Two-way ANOVA with Dunnett’s multiple comparisons test, Fig. 1C). By contrast, MSC and MSC-EV treated DMM mice spent a longer time on the rotarod than untreated DMM mice with no significant difference compared to sham mice at week 16 (DMM+MSCs: 224.8 ± 11.88 sec; p = 0.06, DMM+MSC-EVs: 219.4 ± 20.57 sec; p = 0.09, Two-way ANOVA with Dunnett’s multiple comparisons test, Fig. 1C). Additionally, untreated DMM mice also spend significant less time on the rod at 16 weeks than they did at 4 weeks (week 4: 262.2 ± 10.89 sec, p = 0.0003, unpaired t test, Fig. 1D), while such within group difference was absent in Sham (week 4: 263.7 ± 10.73 sec, p = 0.71, unpaired t test) or treated DMM mice (DMM+MSCs: week 4: 227.7 ± 15.91 sec, p = 0.88, unpaired t test; DMM+MSC-EVs: week 4: 236 ± 14.58 sec, p = 0.3, unpaired t test, Fig. 1D).

We reported previously that mice with joint pain spend less time digging burrows than healthy mice, the digging behavior of mice can thus be considered an ethologically relevant pain assay (*41*). In the digging test, in line with the rotarod test, we observed that untreated DMM mice spend significantly less time digging than sham mice at week 16, while MSC and MSC-EV treated DMM mice exhibit a similar digging duration to sham mice (Sham: 19.79 ± 5.07 sec; DMM: 5.59 ± 2.45 sec, p = 0.03, DMM+MSCs: 15.87 ± 4.59 sec, p = 0.89; DMM+MSCs: 12.32 ± 3.03 sec, p = 0.46; Two-way ANOVA with Dunnett’s multiple comparisons test, Fig. 1E). Consistently, untreated DMM mice dug significantly fewer burrows than sham mice at week 16, whereas the number of burrows dug by MSC and MSC-EV treated DMM mice was similar in number to those dug by sham mice (Sham: 2.77 ± 0.32; DMM: 1.4 ± 0.26; p = 0.02, DMM+MSCs: 2.44 ± 0.53; p = 0.9, DMM+MSCs: 3 ± 0.40; p = 0.95, Two-way ANOVA with Dunnett’s multiple comparisons test, Fig. 1G). However, innate digging differences were observed among mice group. Mice in DMM+MSC-EVs group presented a significantly lower digging duration (week −1: Sham: 26.66 ± 5.16 sec, DMM+MSC-EVs: 4.25 ± 1.51 sec, p = 0.005, Two-way ANOVA with Dunnett’s multiple comparisons test, Fig. 1E) and dug fewer burrows than sham mice pre-surgery (week −1: Sham: 3.55 ± 0.37, DMM+MSC-EVs: 1.37 ± 0.37, p = 0.001, Two-way ANOVA with Dunnett’s multiple comparisons test, Fig. 1G). Comparing to pre-surgery, untreated DMM mice presented reduced digging duration (week −1: 13.84 ± 3.8, p = 0.02, unpaired t test, Fig. 1F) and fewer burrows dug (week −1: 4.11 ± 0.78, p < 0.0001, unpaired t test, Fig. 1H) at 16 weeks, while both sham and MSC treated DMM mice had a similar digging duration (week −1: Sham: 26.66 ± 5.16, p = 0.35, DMM+MSCs: 13.82 ± 2.7, p = 0.7, unpaired t test, Fig. 1F) and number of burrows dug (week −1: Sham: 3.55 ± 0.37, p = 0.13, DMM+MSCs: 2.55 ± 0.47, p = 0.65, unpaired t test, Fig. 1H) as pre-surgery. An increase of both digging duration (week −1: 12.32 ± 3, p = 0.03, unpaired t test, Fig. 1F) and number of burrows dug (week −1: 3.25 ± 0.36, p = 0.003, unpaired t test, Fig. 1H) were seen in MSC+EV treated DMM mice at 16 weeks.

Unlike both the rotarod and digging tests which can only be conducted at set intervals, the DVC^®^system monitors mice activity 24/7. As expected, mice exhibited a high level of activity during the lights off period and compared to the lights on period (Fig. S2A). However, increased irregular activity bouts were seen in DMM mice during the lights on period (i.e. sleep/rest period) in the last week of housing (Fig.S2B, purple box), suggesting a possible rest pattern irregularity in DMM mice caused by pain, similar to the impact of OA on sleep observed in humans (*5*). This irregular activity pattern was computed as regularity disturbance index (RDI), a digital biomarker measuring such irregularity (*42*). We found that DMM mice developed a significantly higher lights on RDI value than sham mice at week 16 (Sham: 0.12 ± 0.028 vs. DMM: 0.45 ± 0.036; p = 0.006, Two-way ANOVA with Dunnett’s multiple comparisons test, Fig. 1I), suggesting a more perturbed rest pattern during lights on in DMM mice. Such an increase in light period RDI was not observed in DMM mice treated with either MSCs or MSC-EVs at week 16 (DMM+MSCs: 0.19 ± 0.036, p = 0.48; DMM+MSC-EVs: 0.23 ± 0.049, p = 0.29; Two-way ANOVA with Dunnett’s multiple comparisons test, Fig. 1I). Similarly, light period RDI of untreated DMM mice at 16 weeks was also significantly higher than their pre-surgery level (week −1: 0.18 ± 0.07, p = 0.002, unpaired t test, Fig. 1J). This rise of RDI was not seen in DMM mice treated with either MSCs (week −1: 0.25 ± 0.04, p = 0.32, unpaired t test, Fig. 1J) or MSC-EVs (week −1: 0.29 ± 0.03, p = 0.36, unpaired t test, Fig. 1J). A decrease of light RDI was observed in Sham mice at 16 weeks (week −1: 0.22 ± 0.01, p = 0.04, unpaired t test, Fig. 1J).

Taken together, these results suggest that MSCs and MSC-EVs both improve pain related behaviors in DMM mice.

**Fig.S1.**
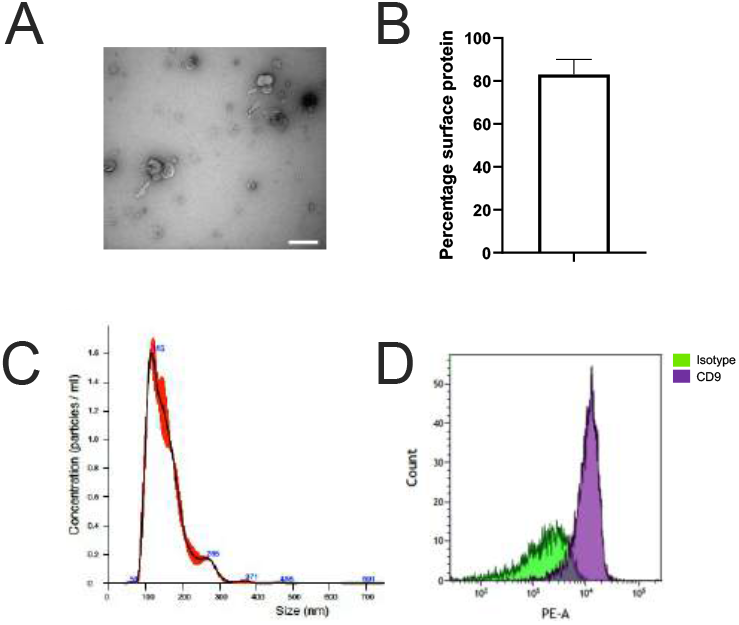
The characterization of MSC-EVs. (A) Representative image of MSC-EVs viewed with a transmission electron microscope, scale bar: 500 nm. (B) Percentage of MSC-EV surface protein. (C) Size distribution of MSC-EVs. Blue numbers indicate the mean particle size at the peak. Red band represent SEM range. (D) Positive signal of surface marker CD9 on MSC-EVs.

**Fig.S2.**
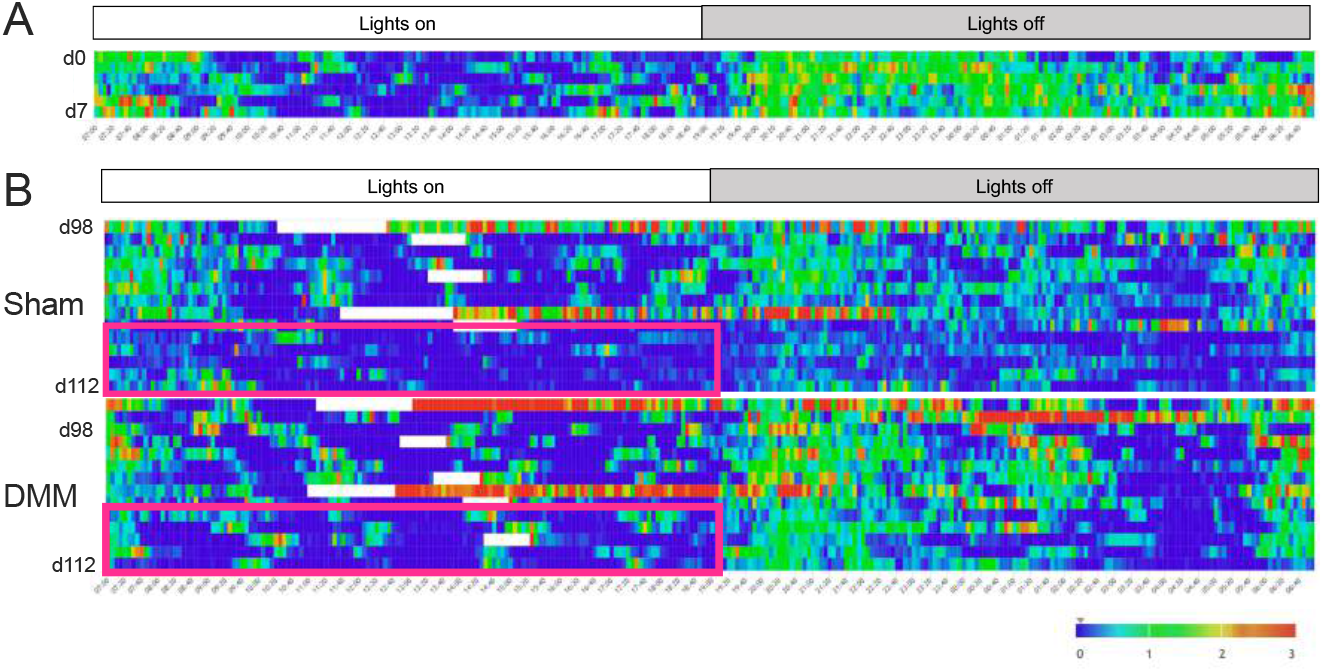
Mouse activity monitored by DVC. (A) Heatmap activity recorded from 3 experimental mice during a week prior than DMM or Sham surgery. Each colored block represents average activities recorded in 5 minutes. The 0-3 scale indicates activity values computed by extruding capacitance change. (B) Heatmap activity of sham and DMM mice from week 14 to week 16 after surgery. d98 and d112 refer to day 98 and day 112 post-surgery. White bars indicate when mice were removed from the cages for experimental procedures or behavioral tests and thus no data were recorded. The purple box shows irregular activity sprouts in DMM mice but not sham mice at week 16. Lights on period: 7:00 – 19:00; Lights off period: 19:00 – 7:00.

### MSCs and MSC-EVs do not improve joint damage in DMM mice

MSCs and MSC-EVs promote cartilage repair in OA joints and have been used as regenerative treatments for OA (*28*). Therefore, we next examined whether the reduction in pain-related behaviors resulted from a lessening of disease progression with regard to joint structure. We performed Safranin O/fast green staining on operated mouse knee joints to evaluate the cartilage damage in different groups and observed that mice from all three DMM operated groups presented with severe joint cartilage damage compared to sham mice (Fig. 2A). We further quantified this observed damage using the Osteoarthritis Research Society International (OARSI) histologic grading system and found that compared to knee joints from sham mice, knee joints from mice in DMM operated groups showed a significantly higher OARSI score on both the medial femoral condyle (MFC) (Sham: 0.39 ± 0.16; DMM: 2.62 ± 0.34; p < 0.0001, DMM+MSCs: 2.24 ± 0.22, p < 0.0001; DMM+MSC-EVs: 3.06 ± 0.35, p < 0.0001; One-way ANOVA with Dunnett’s multiple comparison test, Fig. 2B) and the medial tibial condyle (MTC) (Sham: 0.77 ± 0.16; DMM: 3.08 ± 0.61; p = 0.004, DMM+MSCs: 3.5 ± 0.65, p = 0.003; DMM+MSC-EVs: 3.81 ± 0.6, p = 0.0007; One-way ANOVA with Dunnett’s multiple comparison test, Fig. 2C). These data suggest that MSCs and MSC-EVs do not affect joint damage when injected after 12/14-weeks post-DMM surgery, and that the observed change in pain-related behaviors following MSC/MSC-EV treartment might thus result from an effect of MSCs/MSC-EVs on sensory neurons innervating the knee joint.

**Fig. 2.**
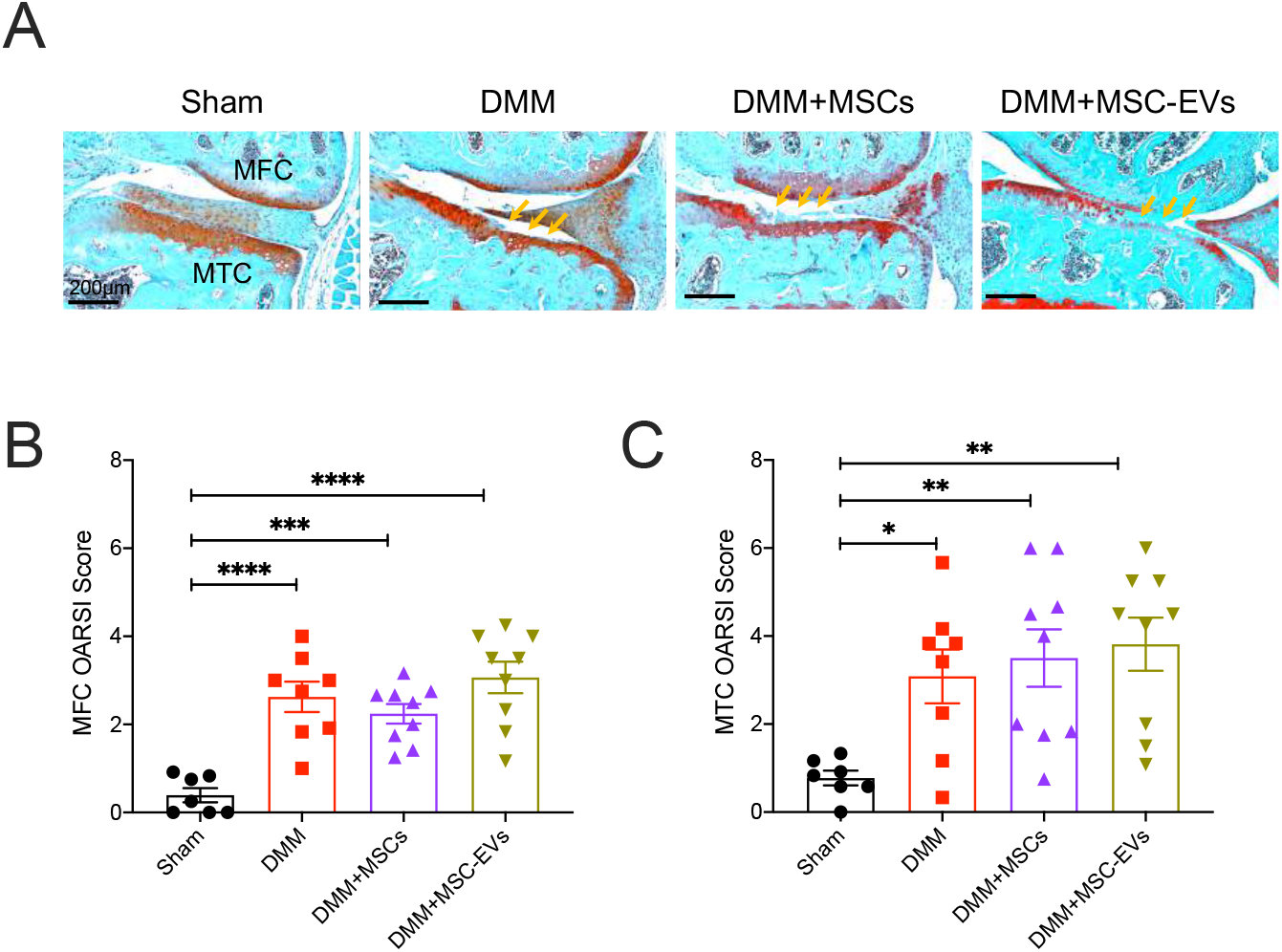
Administration of MSCs or MSC-EVs does not improve knee joint damage in DMM mice. (A) Representative images of Safran O/fast green stained operated knee joint sections from different mouse groups 16 weeks after DMM surgery, scale bar: 200 μm. Cartilage are stained in red. Yellow arrows point cartilage loss (reduced red stain or intact cartilage surface). OARSI score of medial tibia condyle (MTC) (B) and medial femoral condyle (MFC) in different mouse groups. *p<0.05, **p<0.01, ***p<0.001, ****p<0.0001. One-way ANOVA with Dunnett’s multiple comparisons test.

### MSCs and MSC-EVs normalize knee neuron hyperexcitability in DMM mice

We have previously shown that knee-innervating dorsal root ganglion (DRG) sensory neuron excitability increases during acute joint inflammation and that inhibiting function of these neurons normalizes pain-related behaviors (*13*, *41*). In the DMM model, using in vivo Ca^2+^-imaging it has been shown that increased numbers of knee-innervating neurons respond to mechanical stimuli at 8-weeks (*43*), but no in-depth analysis of the excitability of these neurons has been made. Therefore, we injected the retrograde tracer fast blue (FB) into the operated mouse knee joint to label knee-innervating neurons (Fig. 3A). Cell bodies of these labelled neurons were then harvested after mice were sacrificed 16-weeks post-surgery and identified by excitation with a 350 nm light source (Fig. 3A). Using whole cell patch clamp electrophysiology, recording from neurons with similar diameters across groups (Table 1), we found that FB positive neurons in untreated DMM mice have a more depolarized resting membrane potential (RMP) compared to those from sham mice (Sham: −48.96 ± 1.78 mV vs. DMM: −37.52 ± 2.49 mV; p = 0.0009, One-way ANOVA with Dunnett’s multiple comparison test, Fig. 3B) and exhibited a lower action potential (AP) threshold than those knee-innervating neurons from sham mice (Sham: 509.6 ± 45.93 pA vs. DMM: 350.8 ± 37.52 pA; p = 0.03, One-way ANOVA with Dunnett’s multiple comparison test, Fig. 3C), results suggesting that DMM surgery induces knee-innervating neuron hyperexcitability that likely underpins the changes in pain-related behaviors observed. Additionally, the AP of knee-innervating neurons from untreated DMM also had a longer half peak duration (HPD) (Sham: 1.53 ± 0.2 msec vs. DMM: 2.72 ± 0.41 msec; p = 0.019, One-way ANOVA with Dunnett’s multiple comparison test, Fig. 3D) and a longer afterhyperpolarization (AHP) duration (Sham: 17.07 ± 1.38 msec vs. DMM: 29.84 ± 3.54 msec; p = 0.006, One-way ANOVA with Dunnett’s multiple comparison test, Fig. 3F) than knee-innervating neurons from sham mice. When measuring the properties of FB labelled knee-innervating neurons isolated from MSC and MSC-EV treated DMM mice, it was observed that neither their RMP (DMM+MSCs: −44.5 ± 2.03 mV, p = 0.29; DMM+MSC-EVs: −45.25 ± 1.77 mV, p = 0.44, One-way ANOVA with Dunnett’s multiple comparison test, Fig. 3B), nor their AP threshold (DMM+MSCs: 560 ± 43.53 pA, p = 0.71; DMM+MSC-EVs: 607.5 ± 37.79 pA, p = 0.24; One-way ANOVA with Dunnett’s multiple comparison test, Fig. 3C) were significantly different to those of knee-innervating neurons isolated from sham mice, i.e. MSC and MSC-EV treatment normalized DMM induced knee-innervating neuron hyperexcitability. Moreover, the longer HPD and AHP durations seen in knee-innervating neurons isolated from untreated DMM mice were also absent in those neurons isolated from DMM mice treated with MSCs and MSC-EVs (Table 1). As observed AP changes might result from changes in voltage-gated ion channel function, we thus analyzed the properties of macroscopic voltage-gated inward and outward currents (Fig. S3). However, little difference of normalized peak inward current (peak normalized current: Sham: 1 ± 0.08, DMM: 1.05 ± 0.12 p = 0.7, unpaired t test, Fig. S3B) and outward current (peak normalized current: Sham: 1 ± 0.13, DMM: 1 ± 0.1, p = 0.99, unpaired t test, Fig. S3D) was observed among neurons isolated from sham and DMM mice. Thus, data acquired from the other two groups were not analyzed further. Overall, these results suggest that the improved pain-related behavioral change observed in MSC and MSC-EV treated DMM mice results from normalization of knee-innervating neuron hyperexcitability.

**Fig. 3.**
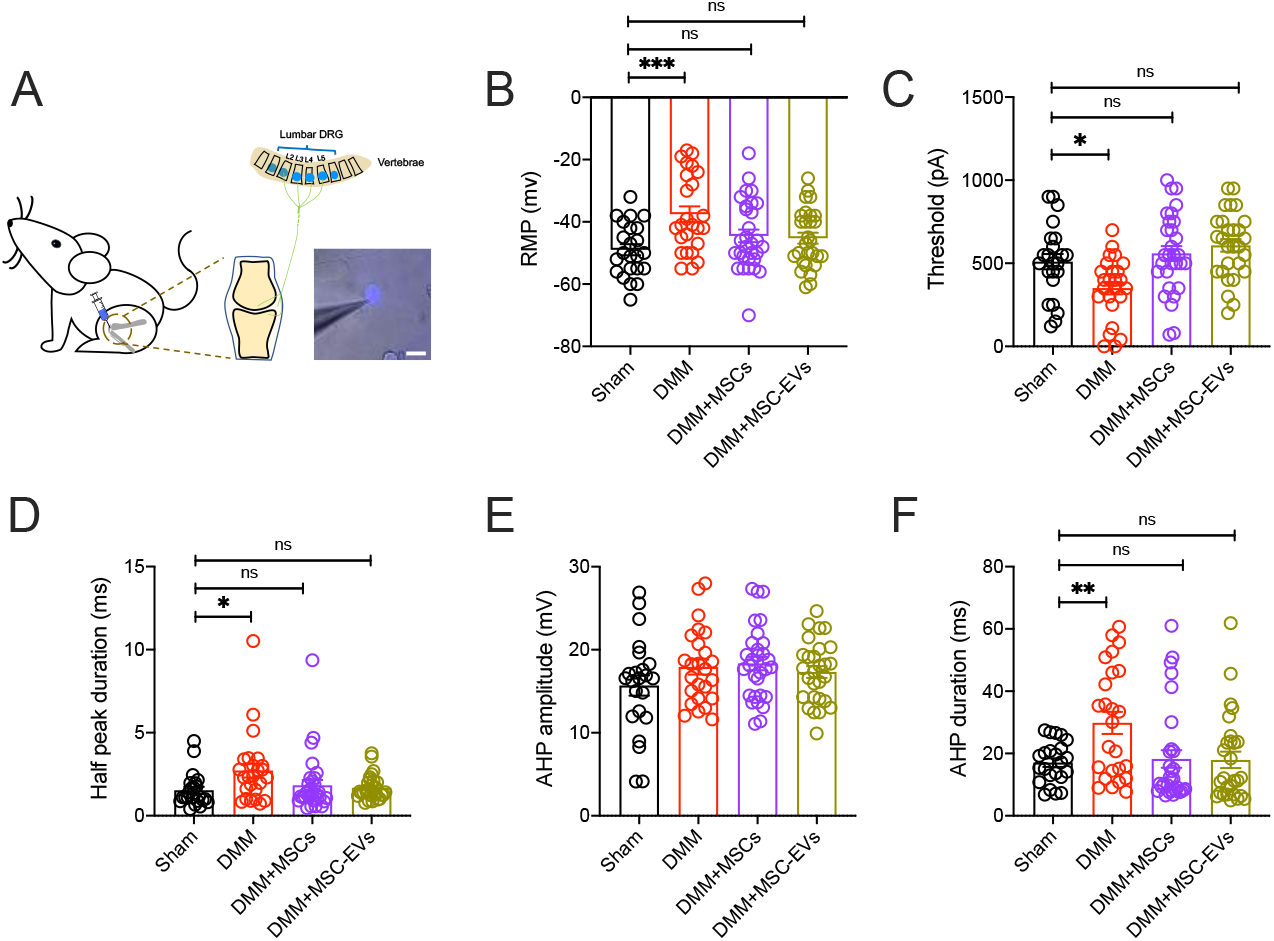
MSCs and MSC-EVs normalize knee-innervating neuron excitability in DMM mice. (A) Retrograde labelling of knee joint innervating neuron by fast blue (FB), scale bar = 50 μm. (B) Resting membrane potential (RMP) of FB labelled DRG neurons isolated from different groups. (C) Threshold of electrical stimulus required for action potential (AP) firing in different FB DRG neurons. AP properties of FB DRG neurons including half peak duration (D), AHP amplitude (E), and AHP duration (F). *p<0.05, **p<0.01, ***p<0.001. ns, no significant difference. One-way ANOVA with Dunnett’s multiple comparisons test.

**Fig.S3.**
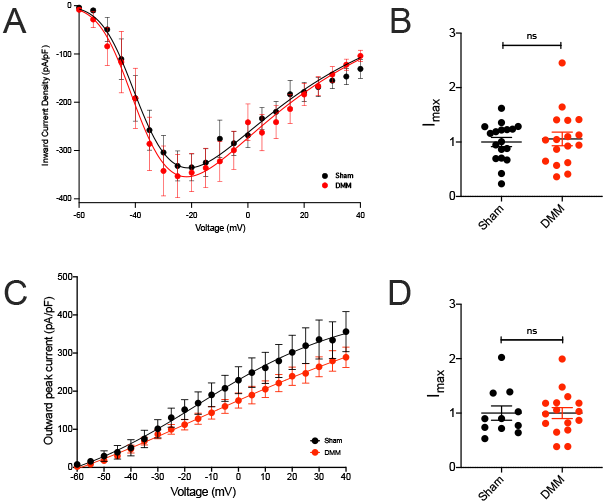
Voltage-gated macroscopic currents of FB neurons. Plots of inward (A) and outward (B) current of FB labelled DRG neurons at different voltage steps normalized by cell capacitance. Peak inward (B) and outward current (D) normalized by maximum current density in sham FB neurons. ns, no significant difference. Unpaired t test.

**Table 1.** Action potential properties of fast blue labelled DRG neurons. RMP = resting membrane potential. n represents neuron numbers; N represents mice number. * signifies p < 0.05 comparing to sham knee neurons, One-way ANOVA with Dunnett’s multiple comparisons test. ^&^ signifies p < 0.05 comparing to DMM knee neurons, One-way ANOVA with Tukey’s post doc test. **,^&&^p<0.01, ***,^&&&^p<0.001.

|  | Sham<br>(n = 23, N = 6) |  | DMM<br>(n = 25, N = 6) |  | DMM+MSCs<br>(n=30, N = 6) |  | DMM+MSC-EVs<br>(n=28, N = 6) |  |
| --- | --- | --- | --- | --- | --- | --- | --- | --- |
|  | Mean | SEM | Mean | SEM | Mean | SEM | Mean | SEM |
| Diameter ($\mu\text{m}$) | 36.01 | 2.17 | 33.2 | 0.79 | 34.03 | 1.00 | 33.91 | 1.08 |
| RMP (mV) | -48.96 | 1.78 | -37.52*** | 2.49 | -44.50& | 2.03 | -45.25& | 1.78 |
| Threshold (pA) | 509.60 | 45.93 | 350.8** | 37.52 | 560&&& | 43.56 | 607.5&&&& | 37.79 |
| Half peak Duration<br>(HPD, ms) | 1.53 | 0.20 | 2.73* | 0.41 | 1.83 | 0.32 | 1.68& | 0.14 |
| Afterhyperpolarization<br>duration (AHP, ms) | 17.07 | 1.38 | 29.84** | 3.54 | 18.27& | 2.82 | 17.93&& | 2.64 |
| Afterhyperpolarization<br>amplitude (AHP, mV) | 15.69 | 1.22 | 17.92 | 0.90 | 18.34 | 0.80 | 17.32 | 0.70 |

### MSC-EVs normalize NGF-induced DRG neuron hyperexcitability *in vitro*

Based on the ability of MSCs and MSC-EVs to induce the same reduction in pain-related behaviors and neuronal hyperexcitability, we hypothesized that the MSC secretome, including MSC-EVs, acts directly upon sensory neurons to normalize their hyperexcitability and in turn reduce pain. Based upon this hypothesis, incubation of DRG sensory neurons with MSC-EVs *in vitro* should be sufficient to normalize neuronal hyperexcitability. To test this hypothesis, we took advantage of the fact that NGF is associated with both OA pain in humans (*44*) and drives pain in the DMM OA model (*45*), as well as directly inducing DRG neuron hyperexcitability *in vitro* (*46*). We established three experimental groups: a Ctrl group with DRG neurons maintained in normal culture medium, an NGF group with DRG neurons incubated with NGF for 40-48-hours, and an NGF + MSC-EVs group in which DRG neurons were incubated in NGF for 24-hours and then NGF + MSC-EVs for 16-24-hours (Fig. 4A). As expected, NGF treated DRG neurons had a lower RMP (Ctrl: −51.78 ± 1.19 mV vs. NGF: −45.48 ± 1.4 mV; p = 0.002, One-way ANOVA with Tukey’s post hoc test, Fig. 4B) and exhibited a lower AP threshold (Ctrl: 706.5 ± 48.22 pA vs. NGF: 568.2 ± 47.39 pA; p = 0.04, One-way ANOVA with Tukey’s post hoc test, Fig. 4C) than the Ctrl group. However, with the addition of MSC-EVs at 24-hours, the RMP of DRG neurons was not significantly different to that of DRG neurons in the Ctrl group (NGF + MSC-EVs: −49.9 ± 1.3 mV, p = 0.059, Oneway ANOVA with Tukey’s post hoc test) and nor was the AP threshold (NGF + MSC-EVs: 730 ± 54.34 pA, p = 0.94, One-way ANOVA with post Tukey test) (Fig. 4B-C). Unlike what was observed in knee-innervating DRG neurons isolated from DMM mice (Fig. 3D,F), no significant change was seen in HPD duration or AHP duration in NGF treated DRG neurons, but in a similar manner to knee-innervating DRG neurons isolated from DMM mice no difference was observed in the AHP amplitude (Fig. 4D-F, summarized in Table 2). We again investigated whether the change in AP threshold might correlate with any change in the properties of voltage-gated ion channel currents. Unlike in knee-innervating neurons isolated from DMM mice, we observed that NGF treated DRG neurons exhibited a larger voltage-gated inward current than Ctrl DRG neurons (peak normalized current: Ctrl: 1.31 ± 0.09, NGF: 2.61 ± 0.42, p = 0.003, One-way ANOVA with Tukey’s post hoc test, Fig. 4G-H) and that this effect was not observed in the NGF + MSC-EV treated DRG neuron group (NGF + MSC-EVs: 1.59 ± 0.27, p = 0.74, One-way ANOVA with Tukey’s post hoc test); no difference was observed in the half-maximal activation potential (V1/2) between Ctrl and NGF neurons (Ctrl: −47.18 ± 1.89, NGF: −50.12 ± 2.15, p = 0.31, unpaired t test). In addition, voltage-gated outward current amplitude was also larger in NGF treated neurons compared to Ctrl DRG neurons, but this was only partially, and not significantly, reversed in neurons from the NGF + MSC-EV treated group (peak normalized current: Ctrl: 1.01 ± 0.08, NGF: 1.81 ± 0.32, p = 0.03, NGF+MSC-EVs: 1.31 ± 0.25, p = 0.6, One-way ANOVA with Tukey’s post hoc test, Fig. 4I-J).

**Fig. 4.**
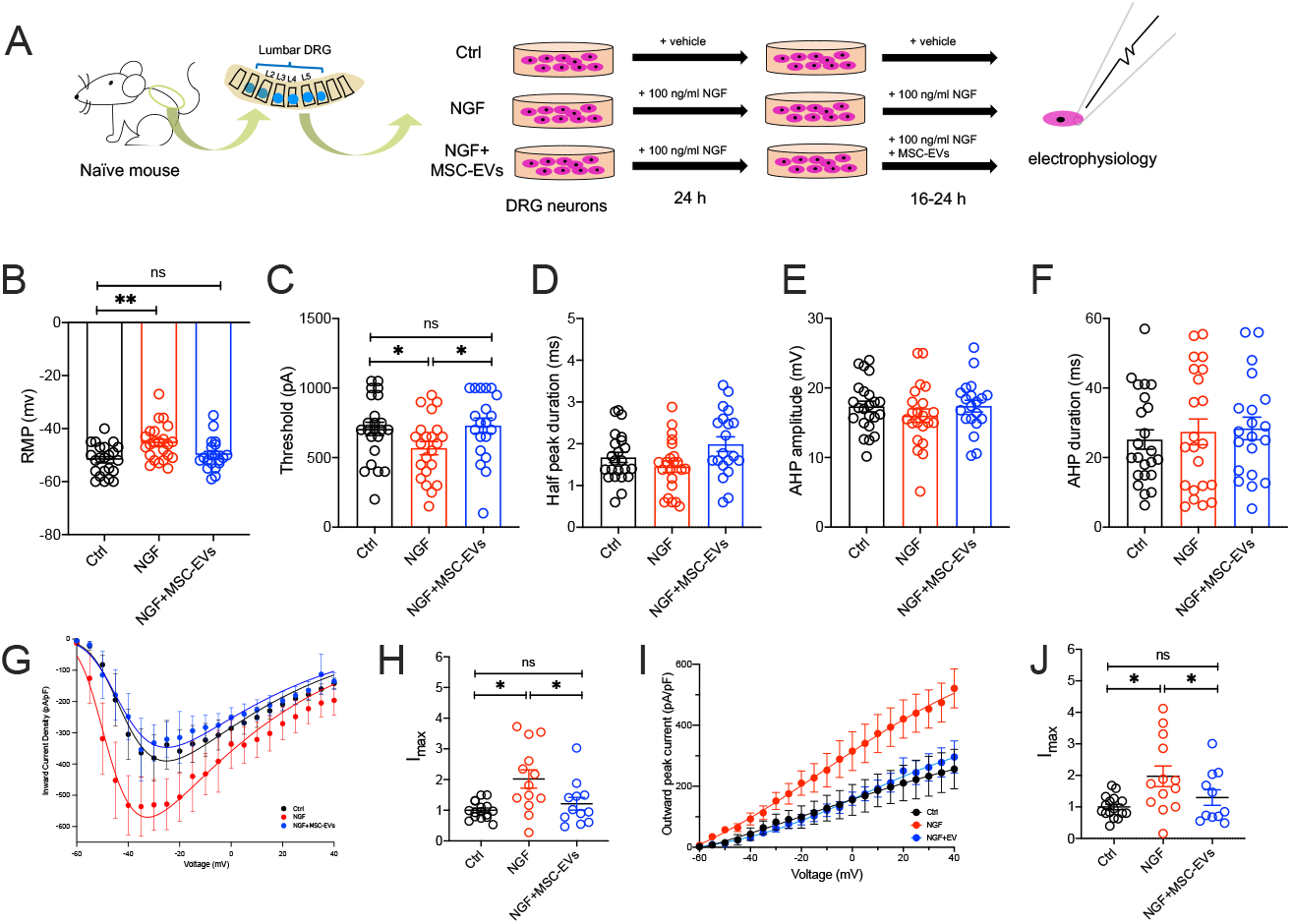
MSC-EVs normalize DRG neuron excitability *in vitro*. ((A) Schematic experimental design of *in vitro* study. (B) RMP of DRG neurons from three different experimental groups. (C) Threshold for AP firing and AP properties including HPD (D), AHP amplitude (E), and AHP duration (F) of DRG neurons from each experimental group. Plots of voltage-gated inward current (G) and outward current (I) density of DRG neurons normalized by cell capacitance in different conditions. Peak voltage-gated inward current (H) and outward current (J) normalized by max current density of Ctrl neurons. *p<0.05, **p<0.01. One-way ANOVA with Tukey’s post doc test.

**Table 2.** Action potential properties of mouse DRG neurons from *in vitro* groups. RMP = resting membrane potential. n represents neuron numbers; N represents mice number. * signifies p < 0.05 comparing to Ctrl group, One-way ANOVA with Dunnett’s multiple comparisons test. ^&^ signifies p < 0.05 comparing to NGF group, One-way ANOVA with Tukey’s post doc tests. **,^&&^p<0.01.

|  | Ctrl (n = 23, N = 4) |  | NGF (n = 23, N = 4) |  | NGF+MSC-EVs (n = 20, N = 4) |  |
| --- | --- | --- | --- | --- | --- | --- |
|  | Mean | SEM | Mean | SEM | Mean | SEM |
| Diameter ($\mu\text{m}$) | 31.51 | 1.01 | 30.93 | 0.78 | 31.24 | 0.95 |
| RMP (mV) | -51.78 | 1.19 | -45.48** | 1.40 | -49.9& | 1.30 |
| Threshold (pA) | 706.5 | 48.22 | 568.2* | 47.39 | 730& | 54.34 |
| Half peak Duration (HPD, ms) | 1.67 | 0.12 | 1.45 | 0.13 | 1.98& | 0.17 |
| Afterhyperpolarization duration (AHP, ms) | 25.18 | 2.77 | 17.39 | 3.7 | 28.38 | 3.25 |
| Afterhyperpolarization amplitude (AHP, mV) | 17.33 | 0.76 | 16.02 | 0.96 | 17.39 | 0.84 |

## Discussion

Numerous pre-clinical (*22*–*24*, *26*–*28*) and clinical studies (*22*) have demonstrated the potential use of MSCs and/or MSC-EVs in treating OA, but the mechanism through which any pain-relieving effects manifest has rarely been examined. When administered at early stages in animal models, both MSCs and MSC-EVs can reduce the extent of disease progression (*27*) and therefore, in this study we deliberately introduced MSCs or MSC-EVs at a time point at which OA and the associated pain behaviors were established to measure if either treatment could specifically ameliorate pain. We found that hyperexcitability of knee-innervating neurons in DMM mice was concomitant with behavior changes and that intra-articular injection of either MSCs or MSC-EVs reduced those same behavior changes, as well as normalizing knee-innervating neuron hyperexcitability. Thus, our results suggest that primary afferent hyperexcitability is causal in DMM OA pain, which supports results of prior studies in rodents and humans showing the importance of primary afferent input in OA pain (*47*), but is the first study to directly measure the excitability of such afferents in the DMM model. MSCs and MSC-EVs have strong immunomodulatory properties and are promising therapeutics for various inflammatory and degenerative diseases, including OA (*31*). While analgesic effects of MSCs are frequently reported in both preclinical and clinical studies (*21*–*24*), mechanisms behind these observations remain elusive. It is recognized that any analgesic effects might originate from immunomodulation and/or chondroprotection, for example, downregulation of inflammatory mediators that sensitize nociceptors in the OA joint (*48*), whereas chondroprotection is perhaps an unlikely mechanism because it has been reported that MSCs reduce pain regardless of regenerative changes in an advanced OA model (*49*). A complication is that OA pain is highly complex with multifactorial mechanisms involved, including both peripheral and central sensitization (*12*). Numerous molecules including NGF, angiotensin-converting enzyme (ACE), and CCL2 have proposed as major drivers of OA pain at the periphery (*18*). Indeed, the blockade of some of these mediators or their receptors produces potent analgesia in OA models (*20*, *50*). MSCs, on the other hand, exert their immunomodulatory effects, at least in part, through inducing overexpression of ACE and CCL2 in inflammatory diseases (*51*–*53*), which might enhance sensitization of knee-innervating sensory neurons leading to pain. Thus, it is possible that undiscovered analgesic mechanisms exist independent of currently known MSC functions.

Consistent with previous analysis (*49*), we observed improved pain-related behavior independent of any regenerative change in OA mouse knee joints following MSC or MSC-EV treatment. In this research, we used three methods to monitor mouse behavior: rotarod, digging assay, and activity monitoring. In the rotarod test, we observed a locomotion deficit in untreated DMM mice at 16 weeks after surgery comparing to Sham mice at 16 weeks post-surgery and to the themselves at 4 weeks post-surgery, consistent with previous reports (*40*, *54*). Such a deficit was not observed in MSC or MSC-EV treated DMM mice. In the digging assay, reduced digging activity was seen in untreated DMM mice, but not Sham or MSC / MSC-EV treated DMM mice at week 16. Undeniably, innate mouse activity difference does exist among mice in different mouse groups. Mice in the DMM+MSC-EVs group had lower digging activity than mice in other groups before surgery, but at 16 weeks the same group presented similar digging activity as mice in the Sham and DMM+MSCs groups, and higher digging activity than their pre-surgery level, which suggests that observed digging difference pre-surgery appears to be compensated by repetitive digging measurements over the 16 weeks experimental period. With activity monitoring, we discovered for the first time that OA mice display enhanced levels of irregular activity during the resting period as disease progresses, similar to sleep disturbances seen in OA patients (50% - 80% of symptomatic OA patients report reduced sleep quality which is positively correlated with pain (*5*, *55*)), while such irregularity was not seen in sham or treated DMM mice at 16 weeks, or in any mice pre-surgery. These results indicate that both MSCs and MSC-EVs normalize the rest pattern in OA mice. Collectively, these data suggested that irregular behavior changes shown in DMM mice were alleviated when DMM mice were treated with either MSCs or MSC-EVs (Fig.1C-J), and such behavior normalization was independent of joint histological improvement (Fig. 2).

Sensory neuron sensitization is known to underlie the pain-related behavioral changes that occur in rat OA (*56*) and sensory neuron hyperexcitability is also common to mouse and sheep models of joint pain (*13*, *41*, *57*). Thus, we performed electrophysiological characterization of retrograde labelled, knee-innervating neurons and observed depolarization of the RMP and lowering of the AP threshold in knee-innervating neurons isolated from DMM mice compared to those isolated from sham mice, effects that were not observed in neurons isolated from DMM mice treated with MSCs or MSC-EVs (Fig. 3). This suggests that normalization of peripheral input may play a role in normalizing behavior. Despite this interesting observation, we acknowledge that normalization of peripheral sensory neuron excitability is unlikely to fully explain the observed behavioral changes as both peripheral and central sensitization components contribute to OA pain, e.g. sensitization of spinal nociceptive reflexes has been observed in a rat OA model (*58*). Whether the improved behavior reported in this study is the result of changes to both peripheral and spinal nociceptive neuron activity change remains unclear. Although the changes observed in primary afferent neuron function could in turn alter spinal circuitry function, it is also possible that spinal circuitry function is also directly influenced by MSC-EVs as these small membrane vesicles are able to pass through the blood-brain barrier and alter neuronal activity in the central nervous system (*59*).

The normalization of peripheral sensory neuron excitability following MSC and MSC-EV injection observed in this study might result from two actions: i) direct action on sensory neurons, and/or ii) reduced nociceptive input/sensitization through modulation of surrounding cellular activity (e.g. reduced release of pro-inflammatory mediators by synoviocytes) (*60*). To address these potential mechanisms, we set up an *in vitro* model to test if MSC-EVs directly alter sensory neuron activity. We induced hypersensitivity in naive mouse DRG neurons by incubating with NGF *in vitro*, which is a major driver of OA pain (*15*) and induces DRG neuron hypersensitivity (*61*). As expected, NGF treated DRG neurons had a depolarized RMP and a lower AP threshold (Fig. 4B-C), which co-incubation with MSC-EVs prevented. This provides initial evidence that MSC-EVs may normalize nociception in the OA joint through direct action on joint sensory neurons, but obviously does not rule out an accompanying indirect effect. However, the NGF treated DRG neurons did not fully recapitulate the changes observed in knee-innervating neurons from DMM mice, e.g. the longer HPD and longer AHP duration seen in knee-innervating neurons isolated from DMM mice were not observed in NGF treated DRG neurons (Fig. 4D, F), and knee-innervating neurons from DMM mice did not exhibit the larger voltage-gated inward currents observed in NGF treated DRG neurons. Consequently, how MSC-EVs modulate neuronal function may differ *in vitro* vs. *in vivo*, but nonetheless data presented here establish models by which the modulatory mechanisms can be further investigated.

Indeed, the molecular mechanisms behind the observed sensory neuron modulation by MSC-EVs remain unknown. Based on current understanding of MSC-EV biology, this phenomenon might be achieved by a variety of different actions. This is because EVs are known to transfer a rich profile of biomolecules (i.e., proteins, lipids, and nucleic acids) to the recipient cells through internalization (*62*). These transferred molecules could alter sensory neuron excitability through modulating ion channel expression or function via different routes. For example, carried microRNAs (e.g. miR-46) can activate second messenger signaling (e.g. p38 MAPK signaling) in neurons and are a key regulator of ion channel activity (*63*), and lipids can act as epigenetic modulators to change ion channel expression (*64*, *65*). Additionally, EVs can also act on cells through direct receptor-ligand binding (*66*), which activates downstream signaling and could lead to changes in ion channel activity. Future research is required to profile MSC-EVs content and identify key molecules influencing sensory neuron excitability in OA pain.

Despite the well-known therapeutic properties of MSCs in OA, their analgesic effects are rarely studied. Our study, for the first time, investigated changes in sensory neuron in the OA joint and how these are altered by the presence of MSCs or MSC-EVs. In doing so, we have discovered that MSC-EVs normalize sensory neuron hyperexcitability both *in vivo* and *in vitro*.This result opens the possibility of using MSC-EVs for chronic pain management and future studies should focus on identifying molecular mechanisms involved in the analgesic effects observed, which raises the possibility of engineering MSC-EVs with enrichment of specific molecules for use as novel pain therapeutics in OA and other chronic pain conditions.

## Material and methods

### Animals

All animal experiments were regulated under the Animals (Scientific Procedures) Act 1986 Amendment Regulations 2012 following ethical review by the University of Cambridge Animal Welfare and Ethical Review Body (AWERB).

A total 36 of C57BL/6J male mice aged between 10 weeks to 12 weeks were used for in vivo study. Mice were purchased from Charles River UK Ltd (Charles River, UK) and assigned into 4 experimental groups of 9 mice: Sham, DMM, DMM+MSCs and DMM+MSC-EVs. All mice were housed in digital individually ventilated cages (DVC) (Cage model GM500, Tecniplast S.p.A., Italy) in a group of 3 with standard water and food supply during the experiment period. Mice were on a normal 12h light/dark cycle at set temperature (21°C) and were regularly monitored by animal technicians, as well as experimenters when undergoing procedures. All the surgical procedures and knee injections performed on mice were carried out under general anesthesia (GA) unless stated otherwise. GA was induced by 4% inhalable isoflurane (Zoties, USA) and maintained by 2.5% (v/v) isoflurane during procedures. Mice were sacrificed after 16 weeks post-surgery by CO2 exposure followed by cervical dislocation.

### Destabilization of the medial meniscus (DMM) surgery

DMM surgery was performed as previously described (*67*). A 3 mm incision was made parallel to the patella on the left leg to expose the stifle joint and the joint capsule was immediately opened using a 15 micro-surgical blade (Swann-Moston, UK). A 30-gauge needle (Terumo AGANI, UK) was used to bluntly dissect the fat pad and expose the medial meniscus (MM). The medial meniscotibial ligament (MMLT) anchoring the medial meniscus to the tibial plateau was carefully cut using a SM65A blade (Swann-Moston, UK). Skin incision was sutured using 6-0 Vicryl^®^ (Ethicon, Belgium). Sham surgery was performed under the same procedure, but without damaging the MMLT. Mice were allowed to recover in a 37 °C chamber (20% oxygen, Tecniplast S.p.A., Italy) with welfare checks every 15 mins for an hour until fully alert and no sign of lameness being present before being returned to their home cages.

### Knee Injections

Stifle injections were performed under general anesthesia using a 10 μl syringe (Hamilton, USA) and a 30-gauge needle (Terumo AGANI, UK) through the patellar tendon. MSCs (2×10^4^ in 6 μl, Lonza, UK) were injected in DMM operated mice at 14 weeks following the surgery. MSC-EVs (6 μl) derived from 2×10^4^ MSCs were injected in to DMM operated mice at 12 weeks and 14 weeks respectively (see supplementary material for MSCs culture, EVs harvest and characterization); MSCs were only injected once as they can continually release mediators, whereas MSC-EVs were injected twice to replenish the supply of mediators. 6 μl of 0.9% saline were injected in untreated DMM and sham mice at 12 and 14 weeks. 1.5 μl retrograde tracer Fast Blue (2% w/v in 0.9% saline; Polysciences, Germany) was injected into the operated stifle joints 7 days prior to mouse sacrifice to label knee innervating neurons.

### Digital ventilated cage (DVC) system

Mice were house in groups of 3 in individual DVC cages with 3 cages in each experimental group. All the DVC cages used are installed on a standard IVC rack (Tecniplast S.p.A., Italy) with external electronic sensors and uniformly distributed 12 contactless electrodes underneath the cage. Animal locomotion activity (referred to as activity in this paper) was monitored by capacitance changes in the electrodes caused by animal movement and computed as previously described (*68*). Weekly rest disturbance index (RDI) during light period was computed to capture irregular animal activity pattern as previously described (*42*). Data was processed and computed on DVC analytic platform (Tecniplast S.p.A., Italy).

### Rotarod

Mouse locomotion and coordination were carried out weekly using a rotarod apparatus (Ugo Basile 47600, Italy) from 4 weeks after surgery (*69*). Mice were placed on the rotarod at constant speed of 4 rmp for 1 min before entering the accelerating testing mode (4 rmp – 40 rmp in 5 mins). Total time spend on the rotarod and the speed at the time of mouse falling, or two passive rotations were recorded. The same protocol was used to train mice one day before the first test.

### Digging

The digging test was carried out weekly in a standard individually ventilated cage (391 × 199 × 160 mm) filled with Aspen midi 8/20 wood chip bedding (LBS Biotechnology) tamped down to a depth of ~4 cm. Each mouse was tested individually in a testing cage for 3 mins without food or water supply after 30 mins habituation in the testing room. Digging training was conducted one day before test. During training, the same digging procedure was carried twice with a 30-mins intermission in-between. All experiments were conducted between 12:30 – 14:30 in the same procedure room and videotaped by a camera (Sony FDR-AX53, UK). Analysis was conducted offline after the conclusion of all studies and following blinding of recordings. Digging duration (time mice spent displacing bedding material using paws) and the number of burrows produced during the testing period was analyzed for all videos by M.A. L.A.P and Q.M. each scored digging duration for a random subset of videos (36% videos were scored by two experimenters, R^2 correlation between scores was 0.95).

### DRG neuron culture

Lumbar DRG (L2-L5) were collected post-mortem and placed into cold dissociation media (L-15 Medium (1×) + GlutaMAX-l (Life Technologies, UK) supplemented with 24 mM NaHCO_3_). Dissected DRG were enzymatically digested in prewarmed collagenase solution (1 mg/ml, 6 mg/ml Bovine serum albumin (BSA) in dissociation media, Sigma, UK) for 15 mins followed trypsin solution (1 mg/ml trypsin, 6 mg/ml Bovine serum albumin (BSA) in dissociation media, Sigma, UK) for 30 mins at 37 °C before mechanical trituration (i.e. pipetting up and down for 8 times). Briefly centrifugation (1000 rmp, 30s) was used to collect neurons from the supernatant. Trituration and centrifugation were repeated for 5 times until 10 ml of supernatant was collected. Collected supernatant was centrifuged at 1000 rmp for 5 mins to obtain cell pellets, which were resuspended in culture media and plated on poly-D-lysine and laminin coated glass bottomed dishes (MatTek, USA). Neurons were incubated at 37 °C, 5% CO_2_ for overnight or 48-hours before electrophysiology depending on the experiments.

### *In vitro* coculture of DRG neurons and MSC-EVs

Lumbar DRG (L2-L5) neurons from non-operated mice (N=4) were isolated and cultured as above, or with addition of mouse nerve growth factor beta (NGF-β, 100 ng/ml). After 24-hours, medium was replaced either without NGF-β, with 100 ng/ml NGF-β, or with NGF plus MSC-EV (10^6^/ml). Neurons were then cultured for another 16-24-hours before electrophysiology recordings.

### Electrophysiology

DRG neurons were bathed in extracellular solution (ECS) (in mM): NaCl (140), KCl (4), CaCl_2_ (2), MgCl_2_ (1), glucose (4), HEPES (10), adjusted to pH 7.4 with NaOH, and osmolarity was adjusted to 280-295 mOsm by sucrose) and recorded by an EPC-10 amplifier (HEKA, Germany) with corresponding software Patchmaster. Patch glass pipettes (4-9 MΩ, Hilgenberg) were pulled by a P-97 Flaming/Brown puller (Sutter Instruments, USA) from borosilicate glass capillaries and loaded with intracellular solution (ICS) (in mM)—KCl (110), NaCl (10), MgCl_2_ (1), EGTA (1), and HEPES (10), adjusted to pH 7.3 with KOH (300-310 mOsm). Ground electrode was placed in the bath to form a closed electric circuit. Fast blue labelled neurons were identified by LED excitation at 365 nm (Cairn Research, UK) with a 450/30× filter tube. Pipette and cell membrane capacitance were compensated by Patchmaster macros and series resistance was compensated by >60%. Resting membrane potential, cell resistance and capacitance were recorded in current-clamp mode. Step current (100 pA to 1000 pA) for 80 ms through 50 steps or no current were injected to generate action potential (AP) under current-clamp mode. AP threshold, half peak duration (HPD, ms), and afterhyperpolarization duration (AHP, ms) and amplitude (mV), were measured in FitMaster (HEKA, Germany) software as previous described (*41*). Voltage-sensitive ion channel activities were assessed under voltage-clamp mode with leak subtraction and series compensation. Cells were held at −120 mV for 240 ms before stepping to the test potential (−60 mV to 50 mV in 5 mV increments) for 40 ms and returned to holding potential (−60 mV) for 200 ms between sweeps. Peak inward and outward voltage-gated current density (pA/pF) were calculated by maximum current (normalized by subtracting average baseline amplitude (5s)) amplitude dividing cell capacitance. Voltage-current relationships were fitted in IgorPro software (Wavemetrics, USA) using the following Boltzmann equation to determine reversal potential (E_rev_) and the half peak activation potential (V_half_):

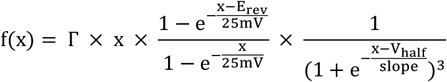

where Γ is the constant, and x is the command potential. To compare the size of current density among neuron groups, the maximum inward or outward current density was normalized to those obtained from the sham neuron with maximum current as Imax.

### Histology

Operated knee joints were collected post-mortem and fixed in 4% (v/v) paraformaldehyde (PFA, Sigma, UK) for 24-hours prior than decalcification. Fixed samples were washed in distilled water for 30 minutes before 21 days of decalcification in 14% (v/v) ethylenediaminetetraacetic acid (EDTA, Sigma, UK) solution (pH 8, adjusted by NaOH pellets) at room temperature (21°C). The completion of decalcification was confirmed through the easy penetration of the tibia bone with a 27G needle. Decalcified joints were processed in graded ethanol series (30, 50, 75, 90, 95, 100 and 100%, 1-hour each), xylene (3×, 1.5-hour each), paraffin (3 ×, 2-hours each) (Fisher, UK) in tissue processor (Leica TP1020 tissue processor, UK) and embedded in paraffin using embedding station (Leica HistoCore Arcadia H embedding station, UK) following routine histological procedures. Embedded samples were sectioned to 7 μm sections using a microtome (Leica RM2235, UK), and mounted on HistoBond slides (StatLab, UK). Slides were deparaffinized and hydrated before staining. Slides were first heated at 60°C for 10 mins following three sequential xylene baths (5mins each), an increased series of ethanol solution (100%, 100%, 95%, 80%, 70%, 50%, 30%; 3 mins each) and distilled water (5 mins) before staining. Hydrated slides were first stained with Weight’s Iron Hematoxylin (Sigma, UK) working solution 7 mins and gently washed with running tap water for 10 mins to remove excessive stain, followed by 3 mins stain with 0.08% (w/v) fast green FCF (Sigma, UK), 10s 1 % (w/v) Acetic acid, and 5 mins 0.1% (w/v) Safranin O (Sigma) before a single dip in 0.5% (w/v) Acetic acid. Slides were then briefly dehydrated with 100% ethanol (2 mins), cleared in xylene (2 mins) and mounted with ProLong^®^ Gold Antifade Mountant (ThermoFisher, UK). Mounted slides were scanned by a PerciPoint O8 microscope and imaged by corresponding ViewPoint software (PerciPoint, Germany). Images were scored blindly by M.A and Q.M using the OARSI scoring system (*70*).

### Statistics

All data are presented as mean ± standard error of mean (SEM). Two-way ANOVA with Dunnett’s multiple comparisons test was used for four groups comparison across time series. One-way ANOVA with Dunnett’s multiple comparisons test was used for four groups comparison with sham group. Unpaired student t-test with was used for two-groups comparisons. Detailed statistical tests are described in individual figure legends. Statistical analysis and graph generation were carried in GraphPad Prism 8.0 software (USA).

## Acknowledgements

Authors acknowledge staff of the Mira Building, University of Cambridge for daily animal maintenance, and MRC Metabolic Diseases Unit, University of Cambridge for tissue processing. Authors thank Karin Newell for assisting animal surgery and substance administration, Dr Toni S. Taylor for the help with digging analysis, and Dr Stefano Gaburro for technical support of DVC analysis.

## Funding

This work was supported by funding from Versus Arthritis (RG21973) to E.S.J.S and Horizon 2020 (RG90905) and Innovate UK (RG87266) to F.M.D.H. W.E.H was supported by the Horizon 2020 (RG90905). L.A.P was supported by the University of Cambridge BBSRC Doctoral Training Program (BB/M011194/1).

## Author contributions

M.A., F.M.D.H., and E.St.J.S. conceptualized the study. M.A. performed the animal surgery, behavior assays, histology, cell culture and electrophysiology experiments, analyzed and visualized data, and draft the manuscript. W.E.H. harvested and characterized extracellular vesicles. L.A.P. performed digging behavior analysis and condition blinding. Q.M. performed digging and histology analysis. F.M.D.H. and E.St.J.S. revised the manuscript. All authors viewed and approved the final form of the manuscript.

## Competing interests

The authors declare no competing interest.

## Data and materials availability

All data needed to evaluate the conclusions in the paper are present in the paper and/or the Supplementary Materials.

## Supplementary Methods

### Extracellular vesicle isolation

Extracellular vesicles were harvested based on previous description (*36*). MSCs were cultured in standard cell culture media α-MEM (Thermo, UK) supplemented with 10% v/v fetal calf serum (thermo, UK), 1% (v/v) Glutamax (100×) (Gibco, UK), 1% (v/v) P/S (Gibco, UK), and incubated at 37 °C, 5% CO2. Passage three MSCs at 80% confluence were switched to serum free culture medium (α-MEM (Thermo, UK), 1% (v/v) Glutamax (100×) (Gibco, UK), 1% (v/v) P/S (Gibco, UK)) for 48-hours incubation. The conditioned medium was then collected and centrifuged at 300 g for 5 minutes, with supernatant transferred to a falcon tube for further centrifugation at 2,000 g for 20 minutes at 4°C. Cell numbers were counted by a hemocytometer. Supernatant was then transferred into polycarbonate ultracentrifuge tubes (Beckman, USA) for differential sequential ultracentrifugation at 10,000 g for 45 minutes and 100,000 g for 90 minutes. Collected pellet was resuspended in PBS for a further ultracentrifugation at 100,000 g for 90 minutes. Newly collected pellet was resuspended in 1ml PBS and stored at −70°C for use.

### Nanoparticle Tracking Analysis

Collected MSC-EVs sample was diluted 1:50 in PBS for Nanoparticle Tracking Analysis (NTA, Malvern, UK). Sample was further diluted from 1:100 to 1:500 with density over 50 particles/frame. Diluted sample was loaded into a NanoSight LM10 Nanoparticle Analysis system following manufacturer’s instruction with a syringe pump rate of 1,000 (Arbitrary units). The analysis was performed in NTA 1.4 analytical software.

### BCA assay

Total surface protein content of MSC-EVs was measured by the Pierce BCA Protein Assay Kit following manufacturer’s instructions (Thermo scientific, UK).

### Transmission electron microscope (TEM)

The MSC-EV suspension was placed on ‘Glow discharge disks’ pre-prepared by the Cambridge Electron Microscopy group. The samples were negatively stained with 2% uranyl acetate in PBS (Sigma, USA) for 2 minutes followed by twice PBS wash and viewed under TEM. Images were acquired by an ORCA HR high resolution CCD camera with a Hamamatsu DCAM board running Image Capture Engine software, version 600.323 (Advanced Microscopy Technology Corp., Danvers, MA, USA).

### Flow cytometry

MSC-EVS were conjugated to 1 μl of 4% aldehyde/sulphate latex beads (Invitrogen, UK) by overnight incubation on a rotary wheel at room temperature with 1ml PBS. 110 μl of 2 M glycine (Sigma, USA) was added following the overnight incubation step (final concentration 200 mM) for 30 minutes before centrifugation at 3,000g for 5 minutes. The sample pellet was resuspended in 1 ml of 0.5% (v/v) FCS in PBS following supernatant removal. Same centrifugation step was applied with pellet was re-suspended in 50 μl of 0.5% (v/v) FCS in PBS afterwards. Resuspended sample was then stained with 1 μl PE anti-human CD9 Antibody (Biologend, UK) at 4 °C for 20 minutes before being diluted in 3ml of 0.5% (v/v) FCS in PBS, centrifuged at 3,000g, and resuspended in 300 μl PBS. Fluorochrome compensation control was prepared by adding one drop of OneComp eBeads (eBioscience, UK) and 0.5 μl of tested antibodies with distinct fluorochrome into 200 μl 0.5% (v/v) FCS in PBS. Prepared samples were stored on ice and scanned by a BD FACS Canto II flow cytometry analyzer (BD Bioscience, UK) within 30 minutes after preparation. Analysis was performed in Kaluza software (Beckman coulter life science, USA) with corrected overlap emission through single stained compensation controls. Only single and live cells were gated during the analysis.

## Notes

### Competing Interest Statement

The authors have declared no competing interest.

